# Chasing the full free energy landscape of neuroreceptor/ligand unbinding by metadynamics simulations

**DOI:** 10.1101/544577

**Authors:** Riccardo Capelli, Anna Bochicchio, GiovanniMaria Piccini, Rodrigo Casasnovas, Paolo Carloni, Michele Parrinello

## Abstract

Predicting the complete free energy landscape associated with protein-ligand unbinding would greatly help designing drugs with highly optimized pharmacokinetics. Here we investigate the unbinding of the iperoxo agonist to its target human neuroreceptor M_2_, embedded in a neuronal membrane. By feeding out-of-equilibrium molecular simulations data in a classification analysis, we identify the few essential reaction coordinates of the process. The full landscape is then reconstructed using an exact enhanced sampling method, well-tempered metadynamics in its funnel variant. The calculations reproduce well the measured affinity, provide a rationale for mutagenesis data and show that the ligand can escape via two different routes. The allosteric modulator LY2119620 turns out to hamper both escapes routes, thus slowing down the unbinding process, as experimentally observed. This computationally affordable protocol is totally general and it can be easily applied to determine the full free energy landscape of membrane receptors/drug interactions.

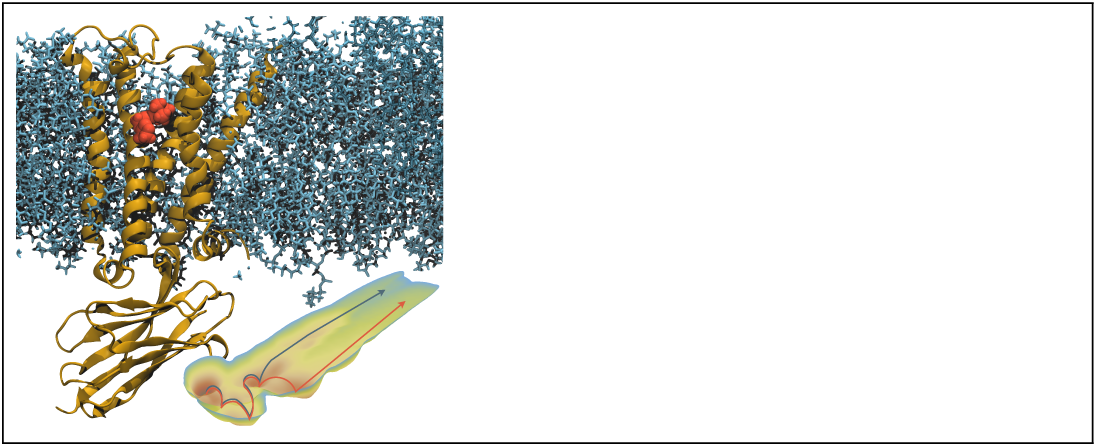
Graphical TOC Entry.

## 1 Introduction

Brain diagnostics rely mostly on Positron Emission Tomography (PET) imaging, where small molecules targeting specific neuroreceptors carry a short-lived isotope.^1^ In their decay process, the isotopes emit a positron whose distribution reveals the position and receptors’ expression levels inside the brain. Neurological diseases (from Schizophrenia to Parkinson’s, from autism to brain tumors^2,3^), can be then readily revealed from receptors maps. A great number of agonist and antagonist that target neuroreceptors for PET, called tracers, has been designed,^4^ yet little is known about their complex binding/unbinding processes. A detailed knowledge of these is of great importance because it allows designing ligands with a more predictable and safer behavior, modulating the affinity and reducing their residence time.^5^

In this work, we investigate *in silico* the molecular recognition process of a widely used tracer, iperoxo (Figure 1), inside the human muscarinic acetylcholine receptor M_2_, a highly valuable target for neuroimaging. The M_2_ receptor is a member of the G-protein coupled receptors (GPCRs) superfamily.^6^ It regulates a large number of different functions in the human body, from modulation of cognition in the brain to motor processes.^7^ It is also involved in neurological and neurodegenerative diseases like schizophrenia and Parkinson’s.^8,9^

**Figure 1:**
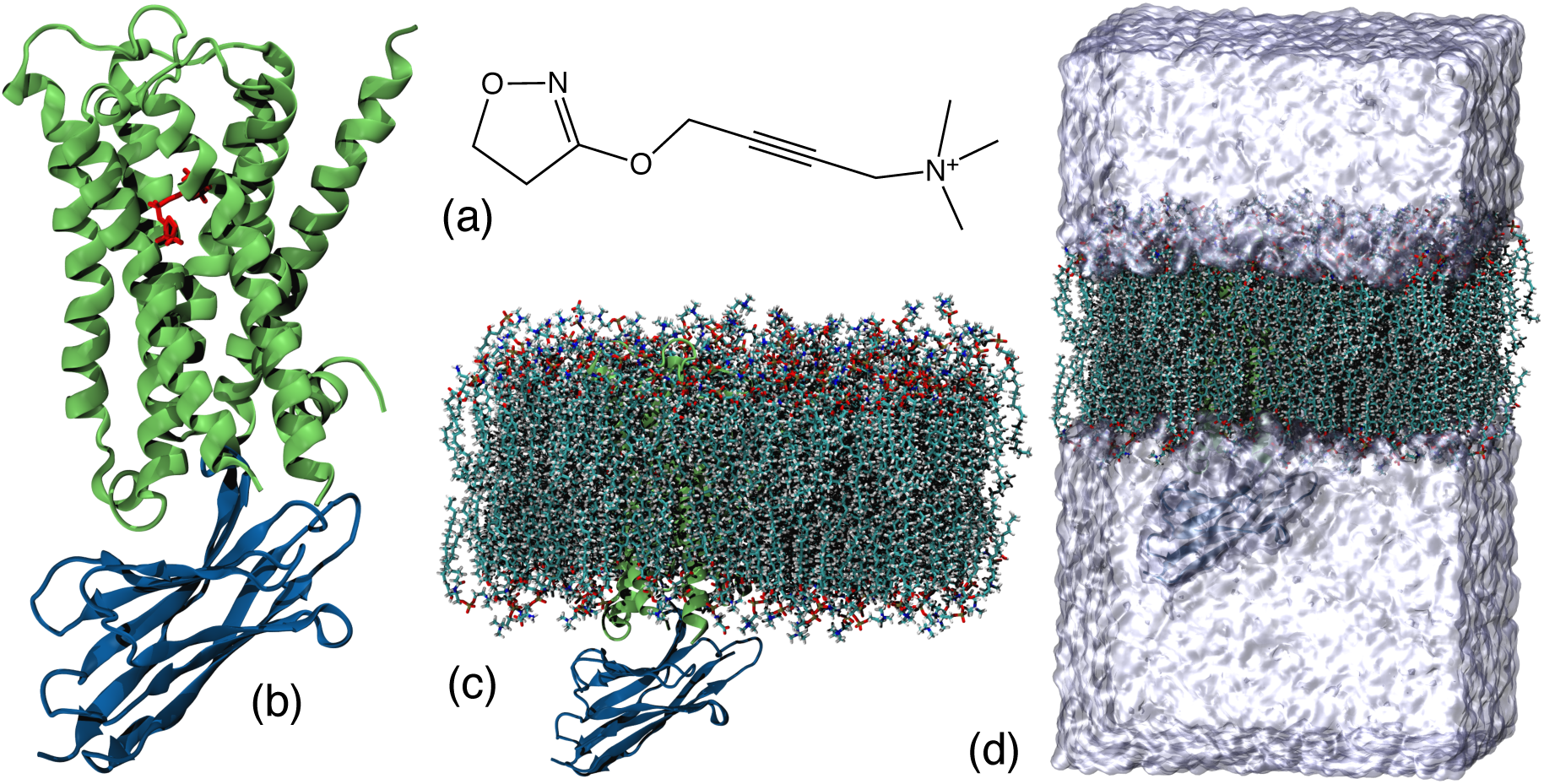
The human M2 receptor in complex with its agonist iperoxo. (a): The ligand (4-(4,5-dihydro-1,2-oxazol-3-yloxy)but-2-ynyl-trimethylazanium) in 2D representation. (b): M2 receptor (green) and its bound ligand, along with the Nb39 nanobody (blue). (c): neuronal membrane with the receptor embedded in it. (d): the simulated system (∼150,000 atoms), formed by ligand, receptor, membrane, water and ions (the latter represented with a transparent surface for sake of clarity).

The description of such complex processes can be tackled on the computer by enhanced sampling techniques based on molecular dynamics simulations.^10^ One of the most used methods is well-tempered metadynamics,^11,12^ where the conformational space of a system can be studied enhancing the fluctuations along some set of relevant functions of the coordinates (called collective variables, CVs), depositing a bias that is dependent on the history of the system. This bias forces the system to explore new states, and, after reaching convergence, it is possible to recover the free energy surface projected along the chosen CVs. This was previously employed to unveil a variety of ligand binding/unbinding processes,^13,14^ moving the problem to a clever identification of a suitable set of collective variables. Efficient metadynamics calculations of binding free energy differences in GPCRs, including M_2_, was recently addressed using a single CV.^15^ However, keeping pharmacological applications in mind, it is highly desirable to use a more general approach in which a larger number of CVs is employed. In this way, the chemical aspects of binding/unbinding transitions and the intermediate state can be better described.^16^

Here, we present a new scheme that leads to a dramatic decrease in the CPU time needed to reach convergence, allowing to use more than one CVs. There is a practical limit in the number of CVs that can be implemented in metadynamics, because the convergence times scales exponentially with the number of the CVs. Considering this point, the number of CVs that is advisable to employ is usually no more than 3, which implies the need of wise dimensionality reduction methods. The problem of dimensionality reduction in complex phase-space transitions has recently seen an increasing number of approaches based on statistical projection of data,^17,18^ as well as methods that involve machine learning/artificial intelligence approaches.^19^ For drug design applications, the first kind of approach appears to be more useful: it keeps the role of every observable employed in the dimensionality reduction in a transparent way, allowing a straightforward interpretation of the final results.

Here we propose to use such approach based on a set of local descriptors, like distances or dihedral angles. These are determined by exploring putative unbinding pathways and candidate intermediate states using the so called Ratchet&Pawl MD (rMD) simulations.^20,21^ These are similar in spirit to steered MD simulations.^22^ Analyzing those states, we were able to extract a set of 23 atom pair distances that describe ligand/receptor non-bonded interactions. The latter were projected in 3 CVs that separate well the intermediate states of the process by means of a recently developed data classification method, called Multi Class Harmonic Linear Discriminant Analysis (MC-HLDA). ^17,18^ This technique highlights the importance of every pair for the landscape description, keeping the physical meaning of the low-dimensional projection evident. Finally, using chemical intuition, we further reduced the number of CVs to 2, dramatically decreasing the converge time.

We use here the funnel variant of well-tempered metadynamics method.^23^ This allows to compute accurate binding free energies at a lower computational cost. The calculations, based on the ligand/protein complex X-ray structure [24], provide us with the full free energy landscape as a function of the two CVs. The resulting binding free energy (−13.6 ± 0.5 kcal/mol) agrees with that measured in saturation binding assays (−13.7 kcal/mol), ^24^ with a convergence time comparable to using a traditional approach with only one CV. Most importantly, our calculations allow for a highly precise description of the ligand binding events. In particular, we were able to confirm the ligand unbinding pathway emerged from rMD analysis and previous metadynamics simulations as a function of one CV,^15^ and to find a second pathway that involves the rearrangement of the extracellular loop 2 (ECL2, Fig. 3).

The study of these pathways provides a rationale for previously reported mutagenesis data,^25^ and, most importantly, an atomic level explanation of the different behavior of the tracer in the presence of an additional allosteric ligand, which enhances tracers affinity for the receptor.^26^ Hence, our method emerges as a very powerful tool for accurate investigations of molecular recognition events of drug binding to human membrane receptors in laboratory realizable conditions.

## 2 Results

Here we describe rMD and metadynamics simulations based on the equilibrated structure of the iperoxo/M_2_ complex in a neuronal membrane mimic environment (See SI). Here, M_2_ is bound to a G-protein mimic nanobody. This allows the receptor to keep the active state conformation. ^24^

### Candidate ligand pathway and intermediate states

Multiple rMD simulations may help identify intermediate states that the ligand finds on its way to unbinding at a moderate computational cost: we simulated 10 different unbinding events (stopping after the ligand is fully solvated) for a total of ∼50 ns worth of simulation. The ratcheting coordinate gives a rough estimation of the progress of the unbinding event. Here, such coordinate is the projection along the direction normal to the membrane of the distance between the center of mass of the ligand and the binding pocket.

By clustering the rMD trajectories, 7 major clusters were identified: the bound state, corresponding to the ligands pose in the X-ray structure complex, ^24^ along with 6 different intermediates. We tested the stability of those states by performing MD simulations that were 50 ns long for every intermediate. Within this time scale only two states (**A** and **C**, Figure 2) remained stable. The others evolved toward directly into state **A** or **C**, or passing through transient state **B** (Figure 2), which was visited several times in different simulations. The picture that seems to emerge from this set of rMD simulations is that the overall process can be described by a rigid rotation of the ligand trimethylammonium group pivoting around D103. Starting from the crystallographic bound state (red in Figure 2), the ligand breaks its non-bonded interactions with hydrophobic residues Q240^(sidechain)^, F195^(backbone)^, V111^(sidechain)^ while rotating around the axis formed by the alkyne bond (see Figure 2). The ligand ring simultaneously forms new H-bonds with Y104^(sidechain)^, S107^(sidechain)^ and Y239^(sidechain)^ (state **A**, dark orange in Figure 2). At this point, the ligand further rotates along the salt bridge (Figure 2), finding a transient state (state **B**, light orange in Figure 2). Here, the ring interacts with Y239^(sidechain)^ and Y262^(sidechain)^. Next, the ligand reaches the last intermediate state (**C**, yellow in Fig. 3). After, it breaks its non-bonded interactions with Y104 while maintaining the H-bonds with Y239^(sidechain)^. The ligand is now partially solvent exposed, forming H-bonds with ∼5 water molecules. Finally, it completely unbinds and it becomes fully solvated. It assumes a conformation which is normal with respect to the membrane. Water molecules could freely go back and forth from the extracellular side to the binding site during the unbinding events. Along with the lipids, they do not play a particular role during the process. The same observations hold in all simulations performed here.

**Figure 2:**
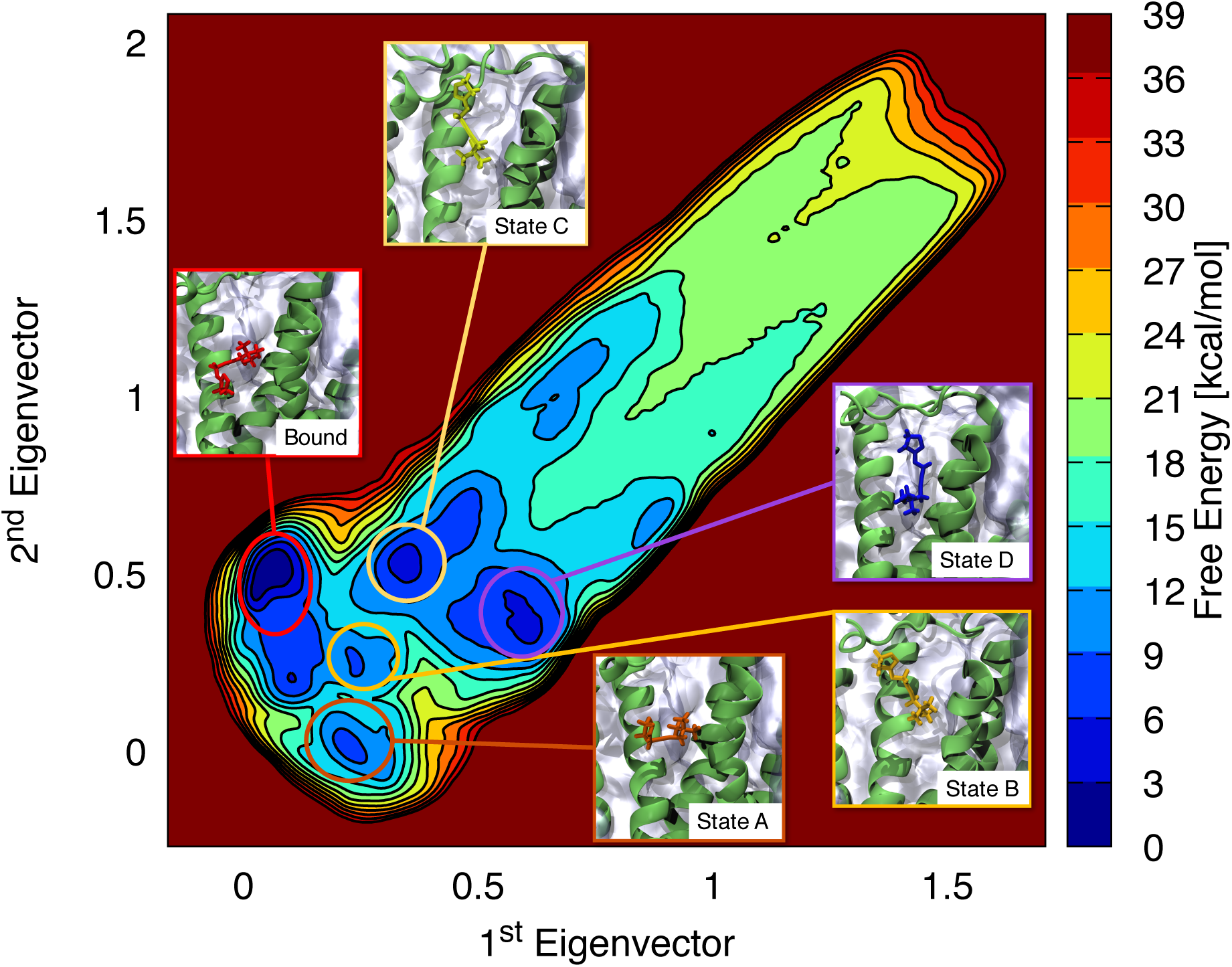
Free energy landscape of ligand unbinding. Free energy surface in function of the first 2 HLDA eigenvectors. The conformations of every free energy minimum of the ligand in the binding pocket are highlighted on the free energy surface.

**Figure 3:**
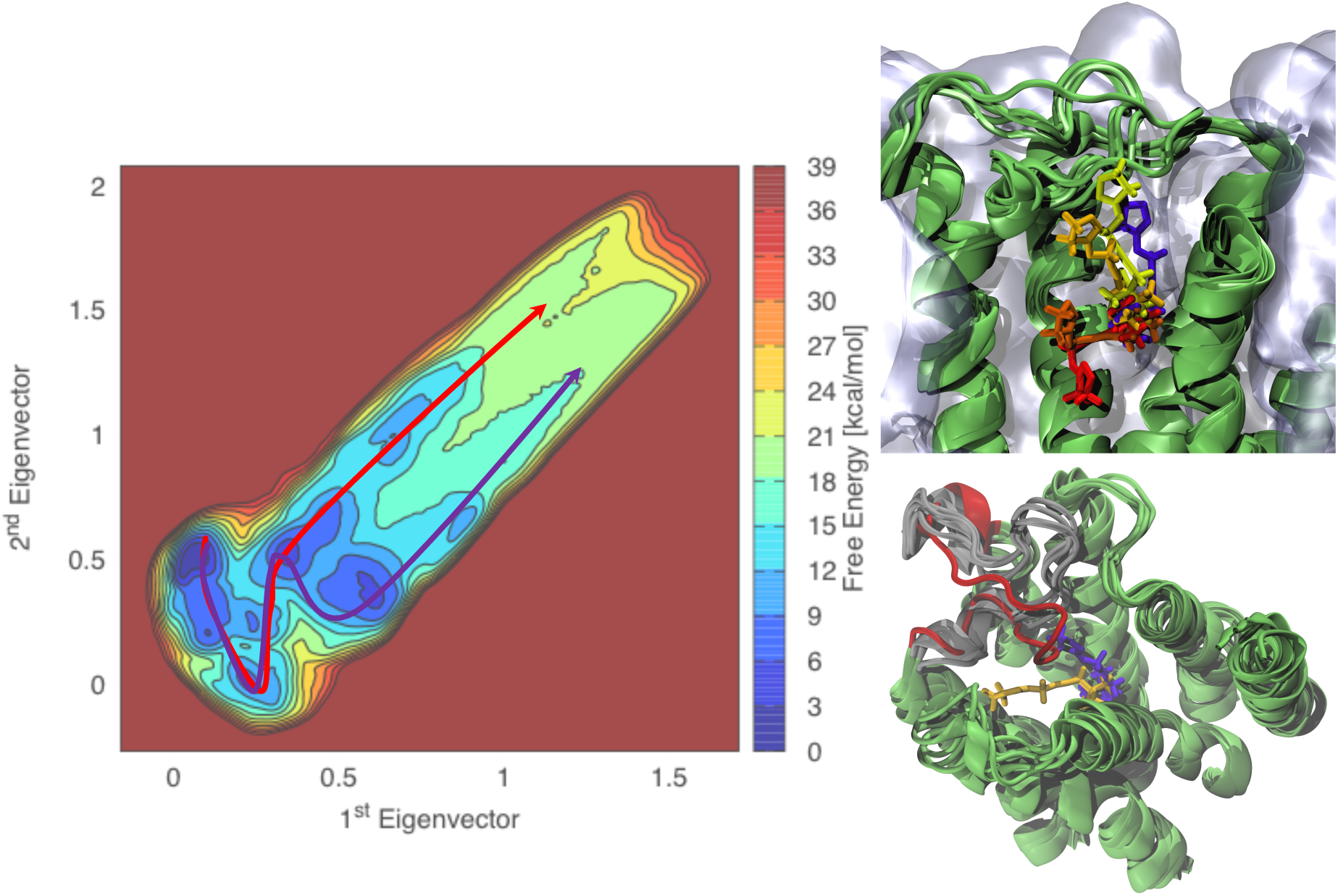
The two pathways of unbinding. Left: schematic representation of the unbinding pathways for iperoxo in the M2 receptor on its free energy surface (Left panel). The first unbinding pathway (red arrow) is the one predicted by exploratory rMD simulations. The second unbinding pathway (violet arrow), is the one found after the rearrangement of the extracellular loop during metadynamics simulations. Top right: Bound (red), states **A** (dark orange), **B** (light orange), **C** (yellow), **D** (violet) for iperoxo (licorice representation) inside the M2 receptor (cartoon representation). Water and ions have been removed for the sake of clarity. Bottom right: cartoon representation of the M2 receptor with the ligand iperoxo in states **C** and **D**. In grey, the position of the ECL2 in the bound state and states **A**, **B**, and **C**. The position of the loop when the system is in state **D** is in red. This change in the receptor conformation forces the ligand to rotate from state **C** (yellow) to state **D** (violet) to reach the solvent.

The simulations allowed to identify 23 local descriptors able to identify all the different intermediate and transient states (see Table S1). These where chosen here as distances between atom pairs involved in the observed non-bonded interaction (i.e. hydrogen bonds and salt bridges) conserved for more than 20% of the simulation time in each state.

In conclusion, the simulations show a rigid rotation of the ligand with the pivot given by the salt bridges between the ligand trimethylammonium group and D103. Consistently, the salt bridge is maintained in all the four states, representing the pivot of the unbinding transition.

### CVs identification

We aim at finding optimal collective variables starting from the 23 descriptors identified as essential transition parameters during the unbiased MD runs describes above. HLDA allows packing into low dimensional yet efficient CVs large sets of state descriptors that capture the essential dynamics of the process. The only information required by the algorithm are simple statistical quantities, namely means and multivariate variances, collected during unbiased runs in the reference states (details in Methods and SI).

To reduce the dimensionality of our descriptors, we exploited HLDA. The key idea of the protocol is to estimate the average and the fluctuations of all the local descriptors (i.e., by computing them in an equilibrium MD simulation) in all the states of interest, and manipulate them in order to obtain a lower dimensional projection of the conformational space that can be used in our enhanced sampling techniques (details in Methods and SI).

In the construction of the HLDA we consider four states, the stable bound state, **A**, and **C**. We added the state **B** because, although it is not stable, it appeared to be an important transit step in the evolution of the unbinding process. Being such state relatively short-lived if compared to states **A** and **C**, it was not possible to simulate it in an equilibrium MD run for enough time. This happens because, during the unbiased MD runs, the ligand jumps to both states **A** and **C**. Therefore, the fluctuation matrix for this state was constructed by combining information coming from all the simulations that explored this state, while for the other 3 states the standard procedure was used (see Methods). According to HLDA, out of these states 3 optimal CVs were constructed. In order to speed up convergence, we neglected the CV with the lowest associated eigenvalue since in our rMD dynamics a strong correlation between the trajectory projection on the lowest and the 2nd highest eigenvalues was seen (see Figure S2).

### Metadynamics simulation

With this choice of CVs, we performed a metadynamics simulation with 8 different walkers. After 1 *μ*s of simulation we have seen a large number of binding/unbinding events in all the walkers and we reached convergence (see Figures S3 and S4). Our metadynamics run shows that all of the rMD intermediate states are relative free energy minima and that the bound state is the absolute minimum (Figure 2). However, another intermediate state (**D** in Figure 2) emerges from our calculated free energy landscape. The ligand here forms H-bonds with Y266^(sidechain)^ and Y104^(sidechain)^, as emerging from a 50 ns-long MD simulation performed on **D**: we have an H-bond with Y266^(sidechain)^ and Y104^(sidechain)^. In addition, ECL2 (Fig. 3) changes its conformation with respect to the bound state: It rotates having as pivot W162 and N183. During this rearrangement, the W90^(backbone)^-T229^(backbone)^ and N246^(sidechain)^-Y177^(sidechain)^ H-bonds are disrupted and the T262^(sidechain)^-C176^(backbone)^, W258^(backbone)^-E175^(sidechain)^, and D173^(sidechain)^-Y83^(sidechain)^ are formed (Instead, in both states a salt bridge between R189 and D97 and a hydrogen bond T247^(sidechain)^-F191^(backbone)^ are maintained). The rearrangement of the extracellular loop forces the ligand to continue the rotation described above having D103 as a pivot to find a way to reach the solvated state.

The presence of intermediate state **D** leads us to suggest the existence of two different pathways from the bound to the solvated state (Figure 3). The first is that identified by rMD (see Section above). In the second one, as we have just seen, the rearrangement of the extracellular loop forces the ligand to perform a further rotation before reaching the solvent (state **D**).

As a last point, we computed the free energy difference between the bound state of our system and the free ligand in the solvent. The free energy difference between the unbound and the bound states obtained was Δ*G* = −13.6±0.5 kcal/mol, in a good agreement with the experimental value of Δ*G* = −13.7 kcal/mol.^24^

## 3 Discussion

We have presented a protocol to study thermodynamics and the intermediate states progression of protein-ligand unbinding processes. As an example, we have focused on a clinically approved tracer binding to a neuroreceptor for PET applications. This is iperoxo in the M_2_ receptor, for which experimental structural information is available. ^24^

To describe the full free energy landscape associated with the process, we have adopted here a two-step strategy. First, we integrated data from out-of-equilibrium technique (rMD) with a modified data classification analysis method (HLDA) to capture the relevant chemical features of the process. Then, with a well-defined and reliable equilibrium method (Funnel Metadynamics) we investigated the thermodynamics of our protein-ligand (un)binding process. The calculations shows that the bound state is the pose in the X-ray structure^24^ and allow identifying the intermediate (**A**, **C**, **D**) and transient (**B**) states (Figure 2).

We calculated relevant observables that can be compared with experimental data, ^24^ obtaining a good agreement (Δ*G*_sim_ = −13.6 0.5 kcal/mol vs. Δ*G*_exp_ = −13.7 kcal/mol). A similar value of Δ*G* was obtained with a single CV. ^15^ However, here we have a much more detailed description and deeper insight of the behavior of the ligand. In particular, Y104 reveals itself as a key residue for the ligand binding process by forming important stabilizing interactions in the intermediates states **A** and **B**. In addition, D103 plays a fundamental role for the conformational changes of iperoxo in the entire unbinding process and in the bound state. Indeed, D103 is the pivot for the rotation of iperoxo (Figure 3 and S7). These results provide a rationale for binding selectivity experiments, which shows that both groups are key residues for agonists binding.^25^ Most importantly, the reconstruction of the full energy landscape points to two different escape pathways for this ligand (Figure 3). The first was identified by rMD simulations and previous computational works.^15^ It does not involve significant changes in the structure of the receptor. In contrast, the second pathway, not emerging from the other simulations, does involve a complex rearrangement of the ECL2 loop (Figure 3), which leads in turn to intramolecular H-bonds breakages.

The presence of allosteric ligands -used to enhanced the ligands affinity^26^-is expected to affect both escape routes. Indeed, visual inspection of the X-ray structure of the receptor in presence not only of the iperoxo tracer but also of one allosteric ligand, called LY2119620 (the only experimental structure available with both the ligands) ^24^ shows that: (i) the first pathway is directly blocked by the allosteric modulator itself (see Figure S6); (ii) the conformational rearrangement of the extracellular loop needed to open the second pathway is hindered (see Figure S6). So, directly or indirectly the allosteric modulator hampers the ligands unbinding from the receptor. This provides a rationale for the large decrease (more than one order of magnitude) in k_off_ upon addition of the allosteric ligand.^27^

Next, we stress the efficiency of our computational protocol: we reached convergence on 2 different CVs with a computational effort comparable to the one needed to converge on the same system on a single CV. ^15^ This is particularly important in large systems (Figure 1) like GPCRs, which constitute a target for more than 30% of FDA drugs.^28^ This protocol can be straightforwardly applied to similar pharmacologically relevant biomolecules, including other transmembrane proteins. Indeed, one can expect that the number of intermediate states in an unbinding transition should be similar to the one found for this system. One can then foresee the possibility of applying the dimensionality reduction given by HLDA to properly describe unbinding events. Keeping pharmacological applications in mind, another important point is the usage of a more realistic neuronal membrane model. The membrane composition was shown to change structure, dynamics and thermodynamics of GPCRs in a significant way.^29,30^

In conclusion, we have shown the possibility to integrate new statistically accurate information towards a more precise drug design. All the non-bonded interaction data obtained by enhanced sampling can give a large number of constraints in current drug design technique, possibly raise the success rate in designing a new molecule that can bind the system studied. In the future, we plan to extend our approach to the calculation of kinetic constants of ligand binding/unbinding, where metadynamics^31^ and other techniques like variationally enhanced sampling^32^ may be exploited.

## 4 Materials and Methods

### System preparation

The structure of the human M_2_/iperoxo complex at 3.5 Å resolution (PDB access code 4MQS^24^) was obtained in the presence of a G-protein-mimic, the Nb_39_ nanobody. The latter stabilizes the receptor in its active conformation. ^24^ We kept the G-protein-mimic in our simulations to avoid that the active state collapses to the inactive one, as observed in the recent MD and accelerated MD simulations.^33^ The crystallographic structure contains some missing loops in the intracellular region that we choose not to reconstruct for the lack of a suitable template (*>*20% sequence identity) among other neuroreceptors. In particular, residue C457 is missing; this cysteine is conserved among a huge number of GPCR and is involved in palmitoylation, which can affect a huge number of function of neuroreceptors. ^34^ In M_2_ receptor, the effect of palmitoylation was experimentally studied, showing that non-palmitoylated cysteine does not show a different affinity for G-protein.^35^ More importantly for our study, it was also shown that the absence of this cysteine does not affect the ligand binding process in the M_2_ muscarinic receptor.^36^

The receptor was inserted into membrane bilayers following the orientation of the OPM database.^37^ The lipids composition was chosen to represent that of a model neuronal membrane^38^ (i.e., 48% cholesterol, 16% phosphatidylcholine (DPPC and POPC), 16% phosphatidylethanolamine (DOPE), 14% sphyngomielin (SM 18:0), 4% phosphatidylserine (DOPS) and 2% phosphatidylinositol (SOPC)). This may be important for the overall fold of the protein and ligand poses.^29^ The system was solvated with explicit water molecules and the appropriate number of sodium and chloride ions were added to neutralize the total charge and reproduce a physiological salt concentration of 150 mM, similar to the one used in affinity experiments.^24^ The final systems contained ∼150.000 atoms and the dimensions of the resulting simulation box was 112×113×149 Å^3^. The dimension along the Z axis was chosen such that complete solvation can be achieved when the ligand dissociates from the M_2_ receptor. The simulations were performed using GROMACS 2016.4. ^39^ The protein, membrane, counterions, water were described by the AMBER ff14SB force field,^40^ the CHARMM-GUI^41^ and Slipids,^42–45^ Joung and Cheatham force fields, ^46^ and TIP3P,^47^ respectively. The force field of the ligand was the generalized AMBER force field (GAFF).^48^ The atomic charges were obtained by the restrained electric potential fitting method (RESP)^49^ with molecular electric potentials obtained in the HF/6-31G* level of theory. Quantum chemical calculations were performed with Gaussian 09. ^50^ A cut-off distance of 12 Å was used for the van der Waals and short-range electrostatic interactions and the long-range electrostatic interactions were computed with the particle-mesh Ewald summation method^51^ using a grid point spacing of 1 Å. Long-range dispersion corrections to the pressure and potential energy were considered.^52^

### MD simulations

We first equilibrated the lipid tails. With all other atoms fixed, the lipid tails were energy minimized for 1000 steps using the steepest descent algorithm and melted with an NVT run for 0.5 ns at 310 K (37°C). In a constant pressure run, we equilibrated the system to a stable volume, fixing only the positions of the protein and the ligand. During this part of the run, that lasted a total of 10 ns, we observed that after 5 ns the volume of the box was fluctuating around its equilibrium value without any canonical drift. We then released the protein restraints for further 0.5 ns keeping the pressure of 1 bar and physiological temperature (i.e. 310 K). After these minimization and equilibration procedures, the production MD simulations were performed on the systems for 0.7 *μ*s. The temperature was kept constant using velocity rescaling thermostat^53^ with solvent, solute and membrane coupled to separate heat baths with coupling constants of 0.5 ps. The pressure was maintained constant with a semi-isotropic scheme, so that the pressure in the membrane plane was controlled separately from the pressure in the membrane normal direction, and the Parrinello-Rahman barostat^54,55^ was applied with a reference pressure of 1 bar, a coupling constant of 2 ps, and a compressibility of 4.5*·*10^−5^ bar^−1^.

### rMD simulations

To study the unbinding path of our ligand we exploited Ratchet&Pawl MD^20,21^ (details in SI). Saleh et al.^15^ used the projection along the direction normal to the membrane of the distance between a conserved tryptophan and the ligand center of mass as CV in their metadynamics simulation. Inspired by this, we defined as our ratcheting coordinate the projection along the direction normal to the membrane of the distance between the heavy atoms of iperoxo ligand and the center of mass of the binding pocket (residues TYR104, SER107, VAL111, PHE195, TYR239). We fixed the bias factor to k=500 kJ/mol/nm and the final ratchet coordinate to rfinal=3 nm. The rMD term was implemented via the PLUMED 2.3 plugin.^56^

### HLDA dimensionality reduction

We made use of the HLDA technique to reduce the number of collective variables from 23 (the number of our local descriptors) to 3 (the number of the bound, intermediate, and transient states minus one). We analyzed the unbiased MD simulation started from all the rMD identified states obtaining the time series of the local descriptors. We then computed averages and standard deviations of the local descriptors in all the states, and then we applied HLDA protocol (details in SI). This dimensionality reduction approach has been used first by Fisher^57^ and adapted to chemical physics problems.^17,18^

### Metadynamics simulations

Here we use a variant of well-tempered metadynamics,^11,12^ called Funnel Metadynamics. ^23^ In this variant, the ligand is free to explore its conformational space inside the binding pocket, but in the proximity of the interface between the solvent and the receptor a funnel-shaped potential is present. The potential confines the ligand in a small region of space outside the receptor. The loss of degrees of freedom given by the funnel potential can be corrected analytically^23,58^ (details in SI). We started the conic region of the funnel potential 13 Å above the crystallographic position of the ligand, reaching the cylindrical part with radius 1 Å to 22 Å above the binding pocket. The cylindrical part of the funnel is 18 Å long to ensure the absence of residual electrostatic interaction of the ligand with the membrane or the receptor at its end (see Figure S5). We performed a Multiple-Walkers ^59^ Well-Tempered Metadynamics, implementing 8 different walkers, each one depositing a gaussian of height kJ/mol every ps with a bias factor of *γ*=24 for a total simulation time of 1 *μ*s (125 ns per replica). Metadynamics was implemented via the PLUMED 2.3 plugin.^56^ All the obtained free energy surfaces were reweighted a posteriori using the algorithm presented by Tiwary and Parrinello.^60^

## Acknowledgement

The authors thank Emiliano Ippoliti, Luca Maggi, and Giulia Rossetti for useful discussion. The authors gratefully acknowledge the computing time granted through JARA-HPC on the supercomputer JURECA at Forschungszentrum Jülich (Project ID: jias58) and acknowledge the JSC for the computing time on the supercomputer JURECA Booster module. This project has received funding from the European Union’s Horizon 2020 Research and Innovation Programme under Grant Agreement No. 785907 (HBP SGA2).

## Supporting Information Available

Details on HLDA, RatchetMD and additional data are available in Supporting Information.

