## Supplemental Information for "Chasing the full free energy landscape of neuroreceptor/ligand unbinding by metadynamics simulations"

### Supplementary Information

#### Supplementary Table 1: Local descriptors of unbinding

| State | ID | atom | residue | iperoxo atom |
|---|---|---|---|---|
| Bound | $d_1$ | ND2 | ASN240 | O2 |
| Bound | $d_2$ | ND2 | ASN240 | N2 |
| Bound | $d_3$ | CB | PHE195 | O2 |
| Bound | $d_4$ | O | ALA194 | C10 |
| Bound | $d_5$ | CE1 | TYR239 | N2 |
| Bound | $d_6$ | CG2 | VAL111 | C10 |
| Bound | $d_7$ | CA | TYR104 | C2 |
| A | $d_8$ | OH | TYR104 | O2 |
| A | $d_9$ | OH | TYR104 | N2 |
| A | $d_{10}$ | OG | SER107 | C1 |
| A | $d_{11}$ | CE1 | TYR104 | C1 |
| A | $d_{12}$ | CD2 | TYR239 | C4 |
| A | $d_{13}$ | CZ | TYR104 | C1 |
| B | $d_{14}$ | OD1 | TYR104 | C1 |
| B | $d_{15}$ | OH | TYR262 | C1 |
| B | $d_{16}$ | CD2 | TYR104 | C7 |
| B | $d_{17}$ | CD1 | TYR239 | N2 |
| C | $d_{18}$ | OH | TYR239 | N2 |
| C | $d_{19}$ | OH | TYR239 | O2 |
| C | $d_{20}$ | CE1 | TYR239 | C8 |
| C | $d_{21}$ | CA | ALA191 | C9 |
| All | $d_{22}$ | OD1 | ASP103 | N1 |
| All | $d_{23}$ | OD2 | ASP103 | N1 |

H-bonds and salt bridges identified in the rMD and persistent in the subsequent 50 ns MD. To be identified as meaningful distances, the H-bonds or salt bridges formed by the atom pairs has to be conserved for more than the 20% of the MD simulation time. The state labeling is the same shown in Figure 2 in the main text. The atom names for the protein residues are the standard ones from Amber14ff, while the names for atoms in iperoxo are explained in Supplementary Figure 10.

### Supplementary Table 2: HLDA dimensionality reduction results

| | Eigenvector | | |
|---|---|---|---|
| | $1^{st}$ | $2^{nd}$ | $3^{rd}$ |
| Eigenvalue | 6487.20 | 2706.27 | 1742.68 |
| $d_1$ | 0.312 | -0.509 | -0.367 |
| $d_2$ | -0.262 | -0.468 | 0.354 |
| $d_3$ | 0.116 | 0.025 | -0.078 |
| $d_4$ | 0.021 | 0.031 | -0.003 |
| $d_5$ | 0.411 | -0.211 | 0.104 |
| $d_6$ | 0.020 | -0.003 | -0.008 |
| $d_7$ | 0.073 | -0.008 | -0.118 |
| $d_8$ | 0.484 | 0.239 | 0.424 |
| $d_9$ | -0.490 | -0.218 | -0.307 |
| $d_{10}$ | 0.004 | -0.002 | -0.011 |
| $d_{11}$ | 0.007 | -0.004 | 0.063 |
| $d_{12}$ | -0.008 | -0.227 | 0.077 |
| $d_{13}$ | -0.002 | -0.019 | -0.059 |
| $d_{14}$ | -0.037 | 0.005 | 0.119 |
| $d_{15}$ | -0.009 | 0.006 | 0.014 |
| $d_{16}$ | -0.001 | -0.006 | 0.004 |
| $d_{17}$ | 0.085 | -0.173 | 0.175 |
| $d_{18}$ | -0.054 | 0.106 | -0.558 |
| $d_{19}$ | -0.347 | 0.430 | 0.237 |
| $d_{20}$ | -0.153 | 0.090 | 0.089 |
| $d_{21}$ | 0.124 | 0.289 | 0.013 |
| $d_{22}$ | 0.021 | 0.015 | 0.021 |
| $d_{23}$ | -0.012 | 0.029 | 0.022 |

Eigenvalues and eigenvectors obtained by our HLDA analysis for the 3 eigenvectors computed here. The vectors are expressed in terms linear combinations of the local descriptors $d_1 - d_{23}$ defined in Supplementary Table 1. The coefficients of these linear combinations are shown here.

### Supplementary Note 1: details on system preparation

The equilibration MD simulation was carried out starting from the X-ray structure of the iperoxo/$M_2$ complex. The system, bound to a G-protein mimetic nanobody and embedded in a model neuronal membrane environment (details in the Methods section), did not show significant deviation from the X-ray structure during the 0.7 $\mu$s-long simulation performed. A notable exception is constituted by the enhanced flexibility of the extracellular loop 2 (Supplementary Figure 1b), which adopts alternative conformations similarly to what observed in the metadynamics simulations (Figure 3). This conformational change is responsible for the "bump" in the root mean square deviation (RMSD) of the protein after 500 ns (Supplementary Figure 1a). When calculated over the receptor helices only, the RMSD approaches an average value of $0.17 \pm 0.02$ nm after only 100 ns, indicating high structural stability. The nanobody as well as the ligand' structures did not deviate from their corresponding crystallographic binding poses (Supplementary Figure 1c). The membrane appears to be well equilibrated and quite ordered as indicated by low average area per lipid (APL) of $0.41 \pm 0.01$ nm$^2$ (Supplementary Figure 1d). This value is consistent with other MD and NMR studies [8, 7] of model membranes characterized by a high cholesterol content (in our case 50% of the total lipids composition).

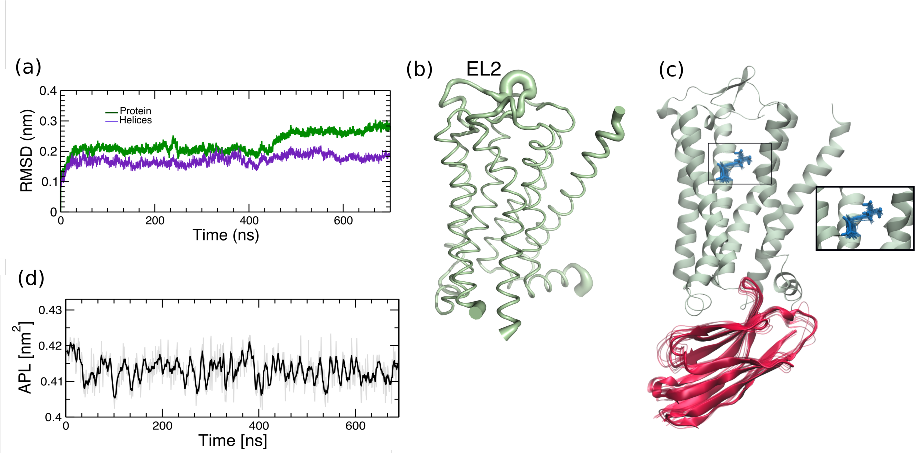


Supplementary Figure 1: (a) RMSD with respect to the protein X-ray structure calculated for the entire protein (green line) and for the protein helices (violet). (b) Protein root mean square fluctuations (RMSF) during the simulation. RMSF values are represented as putty cartoon: ticker areas correspond to highly fluctuating residues. (c) Selected structures of the nanobody and of iperoxo ligand during simulation. (d) Area per lipid (APL) over time calculated with the g_lomepro software [6] on a 100 × 100 grid.

### Supplementary Note 2: details of Ratchet&Pawl MD

Ratcheted Molecular Dynamics (rMD)[4, 5] is a non-equilibrium sampling technique. This method allows us to explore the transition path of a system between two points. Knowing the starting and the end point of transition of interest, a scalar ratcheting coordinate $r(t)$ that connects those points (reaching the ratcheting coordinate $r_{\text{final}}$ that corresponds to the end point) has to be defined. At this point, a bias that dumps thermal fluctuations in the direction opposite to the end point is applied. This is achieved by imposing a ratchet potential:

$$V_{\text{rMD}}(\xi(t)) = \begin{cases} \frac{k}{2}\left(\xi(t) - \xi_m\right)^2 & \xi(t) > \xi_m(t) \\ 0 & \xi(t) \le \xi_m(t) \end{cases} \tag{1}$$

where

$$\xi(t) = (r(t) - r_{\text{final}})^2$$

and

$$\xi_m(t) = \max_{t \in [0,T]} \xi(t)$$

### Supplementary Note 3: details of HLDA technique

We have a system that displays a certain number $k = 1, ..., K$ of states and a number $i = 1, ..., N_d$ of descriptors $d_i(\mathbf{R})$. For each of the $K$ states we calculate the expectation value $\boldsymbol{\mu}_i$ of every $d_i(\mathbf{R})$ and their correlation matrix $\boldsymbol{\Sigma}_i$. These can be computed implementing data that come from MD runs that explore the local conformational space of each state. We then search for the linear projection into a $(K-1)$-dimensional space that best separates the states. The rectangular matrix $\mathbf{W}$ of dimension $(K-1) \times N_d$ that achieve this result is obtained by maximizing the ratio

$$\mathcal{J}(\mathbf{W}) = \frac{\mathbf{W}^T \mathbf{S}_b \mathbf{W}}{\mathbf{W}^T \mathbf{S}_w \mathbf{W}} \tag{2}$$

where the scatter matrices $\mathbf{S}_b$ and $\mathbf{S}_w$ are defined as

$$\mathbf{S}_b = \sum_{k=1}^{K} (\boldsymbol{\mu}_k - \overline{\boldsymbol{\mu}})(\overline{\boldsymbol{\mu}} - \boldsymbol{\mu}_k)^T \tag{3}$$

$$\mathbf{S}_w = \sum_{k=1}^{K} \frac{1}{\frac{1}{\boldsymbol{\Sigma}_k}} \tag{4}$$

where $\overline{\boldsymbol{\mu}}$ is the overall mean of all the datasets.
To maximize the ratio in eq.(2), we can solve the generalized eigenvalue problem

$$\mathbf{S}_b \mathbf{W} = \lambda \mathbf{S}_w \mathbf{W} \tag{5}$$

that, assuming $\mathbf{S}_w$ invertible, gives us the standard eigenvalues problem

$$\mathbf{S}_w^{-1} \mathbf{S}_b \mathbf{W} = \lambda \mathbf{W} \tag{6}$$

We will get $K-1$ non-zero eigenvalues, where the eigenvector are the coefficients of our linear combination that defines the axes of our projection. The magnitude of the eigenvalues can be interpreted as an approximate estimation of the importance of that CV in states discrimination.
This dimensionality reduction approach has been used first by Fisher [1] defining $\mathbf{S}_w$ as an arithmetical average, and later adapted to chemical physics problems by defining $\mathbf{S}_w$ as the harmonic average.

### Supplementary Note 4: Entropic correction for Funnel Metadynamics

During a metadynamics calculation in its Funnel variant, the molecule is restrained to remain in a funnel-shaped potential that limits its possibility to freely explore the environment in the solvent. This constrining favors the recrossing and rebinding events, speeding up convergence. At the same time, an entropy loss is observed. This missing entropic contribution can be estimated analytically [2, 3] to obtain the absolute protein-ligand binding free energy $\Delta G_b^0$ as

$$\Delta G_b^0 = \Delta G - k_B T \ln\left(\pi R_{\text{cyl}}^2 C^0\right) \tag{7}$$

where $\Delta G$ is the free energy difference obtained by metadynamics calculation, $k_B$ is the Boltzmann constant, $T$ is the system temperature, $R_{\text{cyl}}$ is the width of the funnel cylinder, and $C^0$ is the standard concentration.

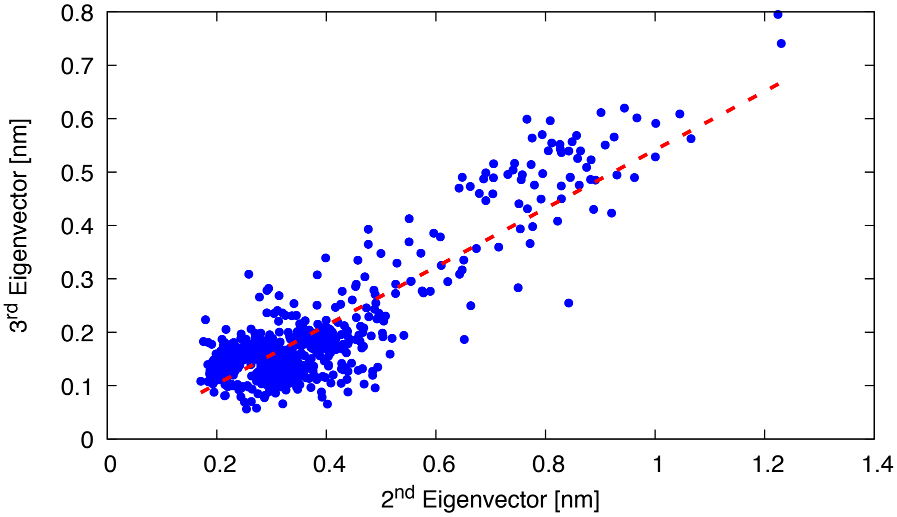


Supplementary Figure 2: Correlation between $2^{\text{nd}}$ and the $3^{\text{rd}}$ HLDA eigenvectors. Scatterplot of the projection of the rMD unbinding trajectories along the $2^{\text{nd}}$ and the $3^{\text{rd}}$ eigenvectors. The two projections are strongly correlated ($r = 0.874$). The redundancy of the information contained in the two eigenvectors convinced us to reject the one with the lower eigenvalue in our metadynamics calculations.

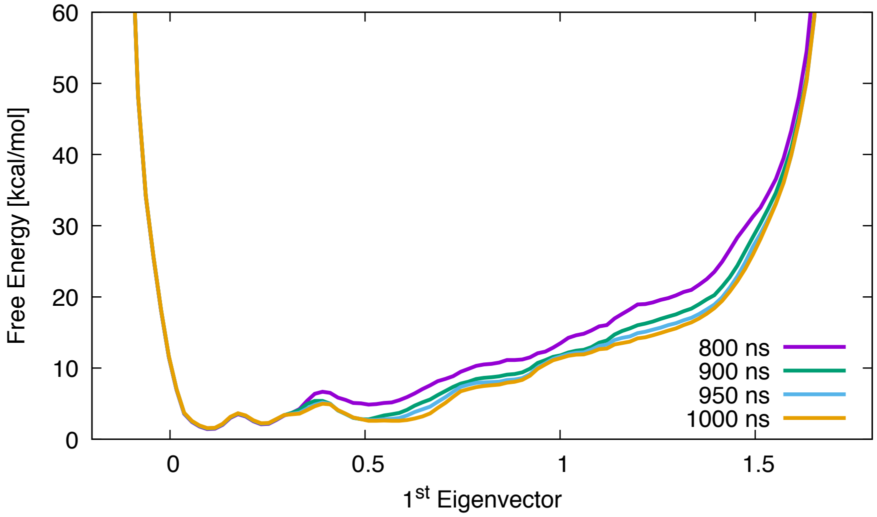


Supplementary Figure 3: Convergence of the free energy surface along the first HLDA eigenvector. We can see that after 900 ns the bound and the unbound state were already explored by metadynamics. The similarity between the free energy surfaces after 950 ns and 1 $\mu$s suggests that we reached convergence. The free energy profile does not show the second entropic minimum (unbound state) in the free energy surface because the funnel potential attenuates the entropic effect.

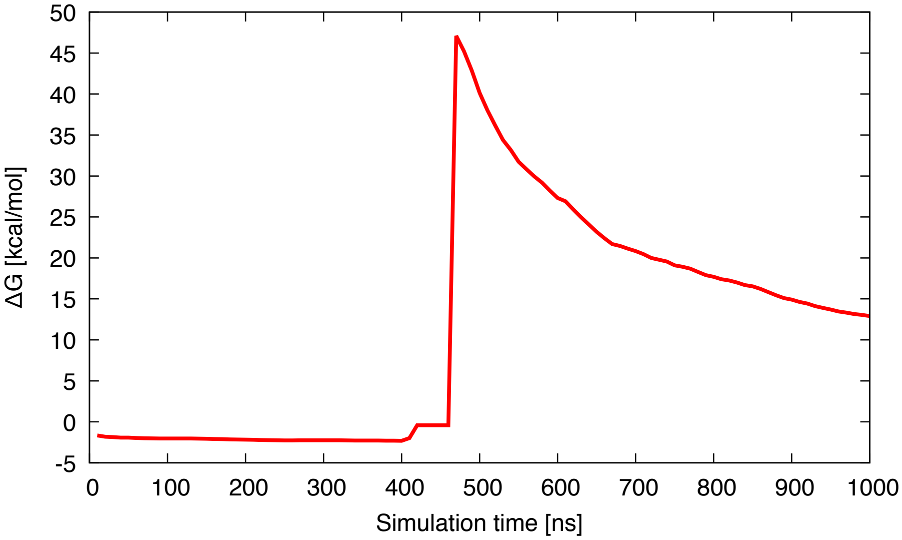


Supplementary Figure 4: Free energy difference value between the bound and the unbound state as a function of time. At the beginning (between 0 and 450 ns), the walkers explored the region inside the binding pocket. After the first unbinding transitions ($\sim$ 460 ns), the bias is deposited also in the region outside the receptor, converging slowly to a stedy value.

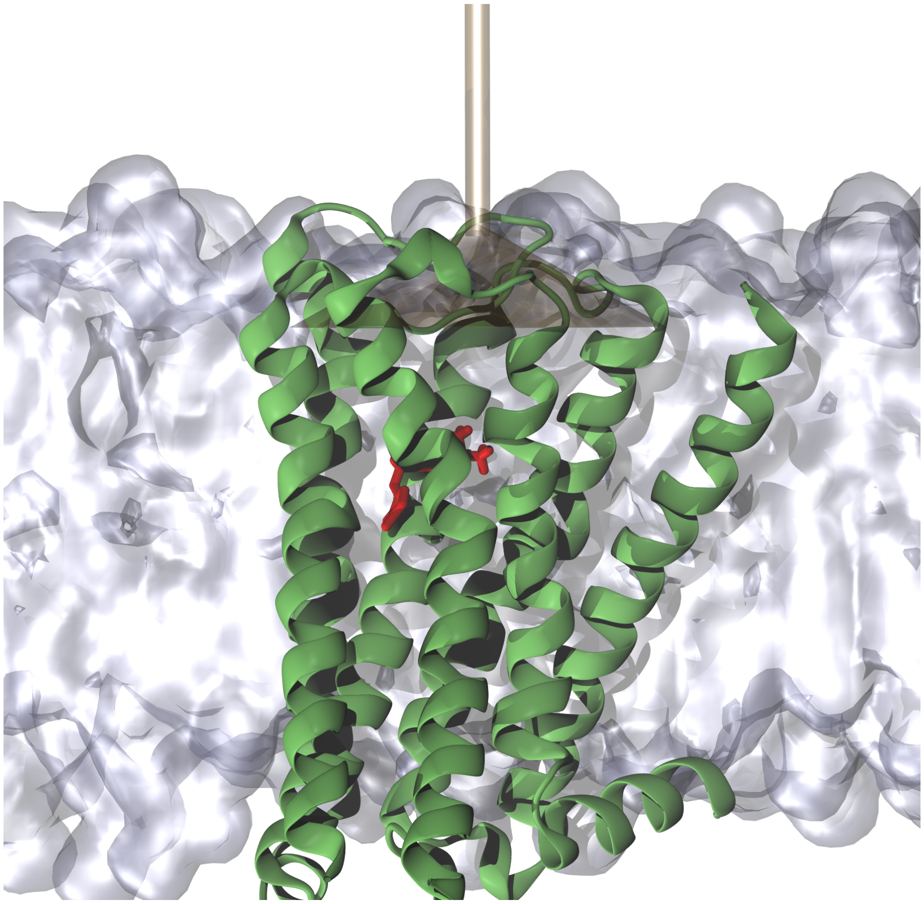

Supplementary Figure 5: Restraining potential position in Funnel metadynamics simulation. Here we show the position of the funnel potential (gold) with respect to the receptor (green), the ligand (red), and the membrane (grey surface). Water and ions are removed for sake of clarity.

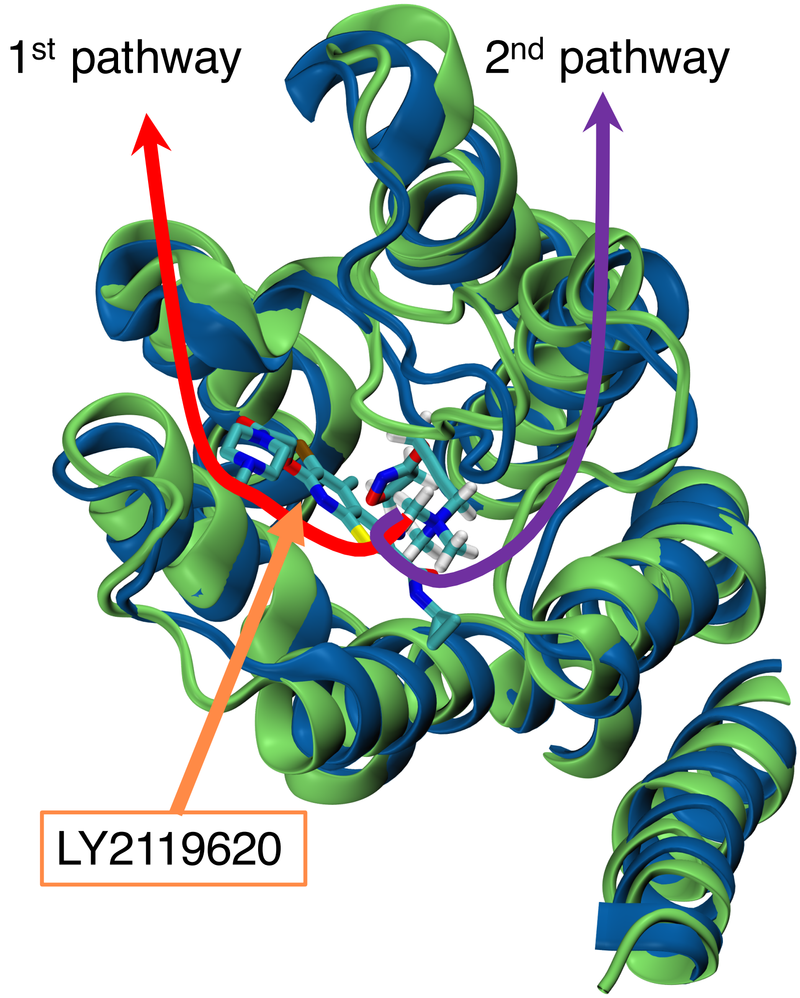


Supplementary Figure 6: Pose of the iperoxo ligand in the bound state and in ternary complex with the allosteric ligand LY2119620. The figure has been obtained by superimposing the bound state (green) with the crystallographic structure of the $M_2$ receptor+iperoxo+LY2119620 (blue). LY2119620 is located in the middle of the first pathway (red arrow), and also prevent the rearrangement of the ECL2 to open the second pathway (violet arrow).

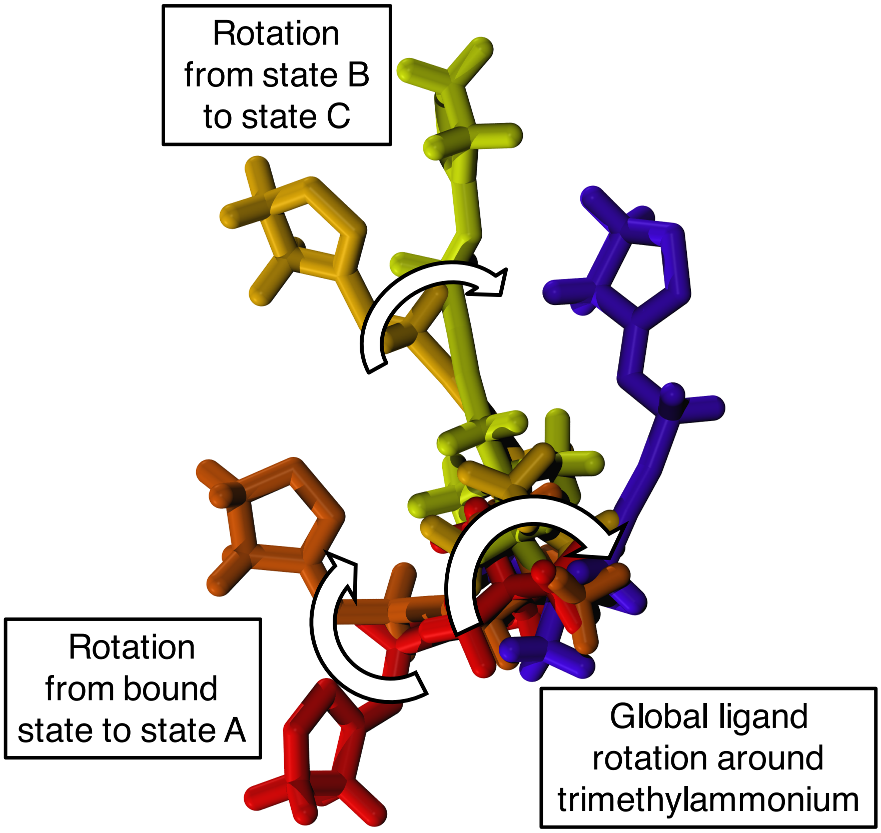


Supplementary Figure 7: Unbinding process for the ligand iperoxo. As detailed in the main text, we see an initial rotation around the alkynic group (atoms C2-C3 in Supplementary Figure 9) from bound state (red) to state A (dark orange), followed by the global rotation around the trimetylammonium group (atoms C5, C6, C7, N1, C1 in Supplementary Figure 9) that make the ligand reach the state B (light orange), then state C (in yellow, here performing another local rotation as detailed in figure) and state D (violet). The state labeling follows the same labeling in the main text (see also Figure 2 in main text).

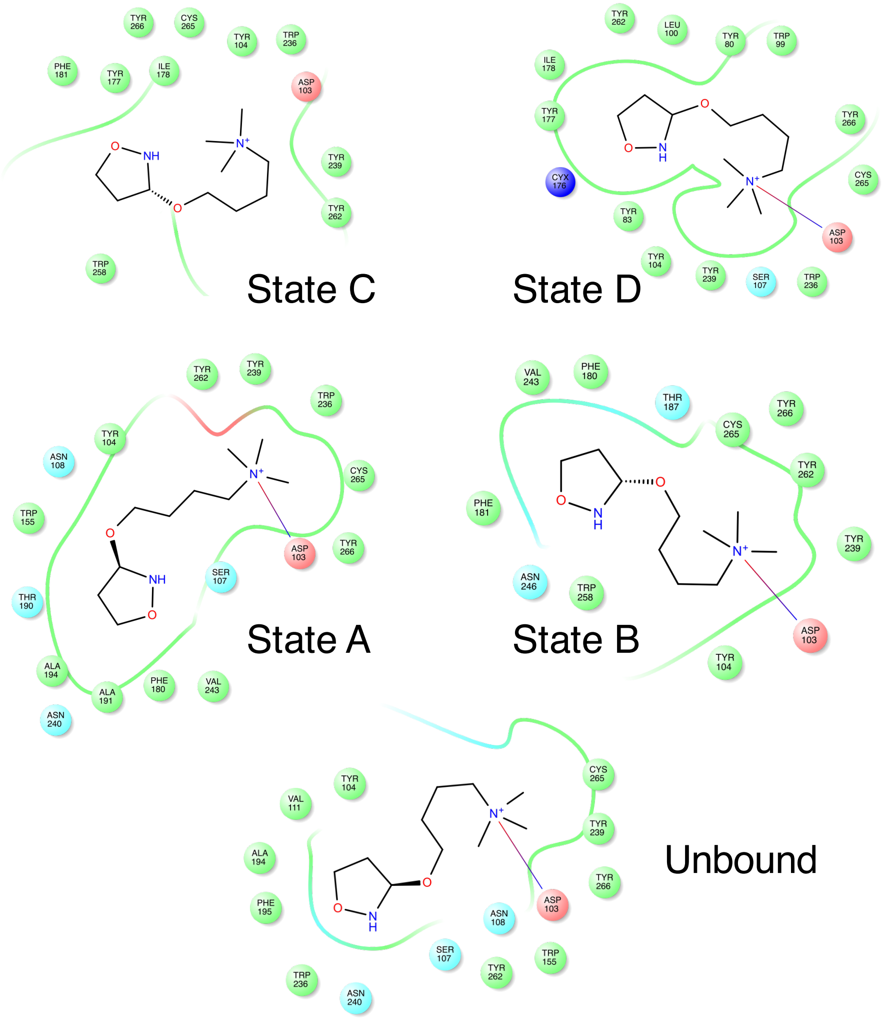


Supplementary Figure 8: $M_2$ receptor/iperoxo interactions in the 5 states of Figure 2. We sketched the ligand surrounded by the residues within 3.5 Å.

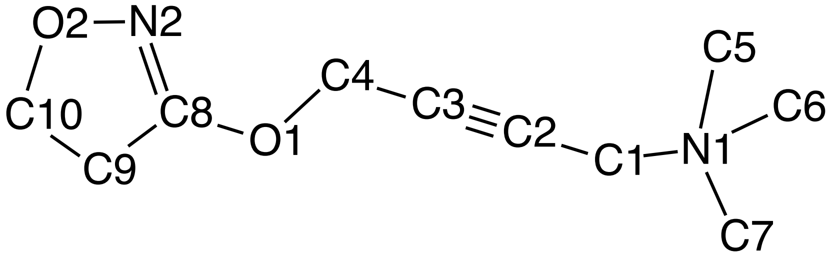


Supplementary Figure 9: Iperoxo with non-hydrogen atom names used in our force field.

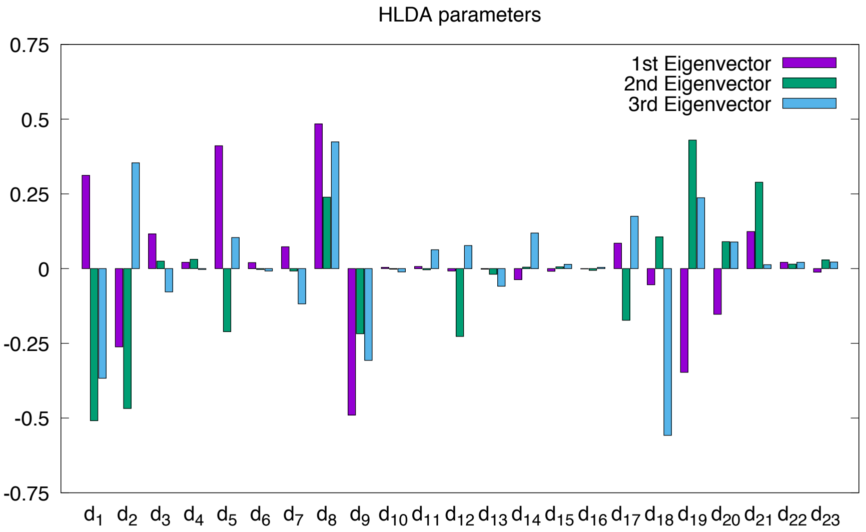


Supplementary Figure 10: Histogram with the weight of the local descriptor in every eigenvector obtained by HLDA.
